# Laboratory evolution of *Mycobacterium smegmatis* in the presence of a fluoroquinolone leads to extreme drug resistance phenotype through the over-expression of *Msmeg_5659-61 efflux pump.*

**DOI:** 10.1101/2022.12.18.519879

**Authors:** Deepika Rai, Priyanka Padwal, Priyanka Purkayastha, Sarika Mehra

## Abstract

Resistance to multiple drugs is one of the significant barriers in the treatment of tuberculosis (TB). Knowledge of mechanisms of resistance is important to design effective treatment strategies. While mutations in genes coding for drug targets are thought to be the primary source of drug resistance, absence of mutations in these genes in many clinical strains suggests additional mechanisms of resistance. In this study, we employ adaptive laboratory evolution of *Mycobacterium smegmatis* to understand alternate mechanisms of drug resistance to norfloxacin, a fluoroquinolone (FQ). Results show that, in addition to fluoroquinolones, the evolved strain, Nor^r^, is resistant to first-line drugs, rifampicin and isoniazid, and a second-line drug (amikacin), exhibiting extreme drug resistance phenotype. However, mutations were absent in any of the drug target genes. Drug uptake studies revealed that resistance is an attribute of decreased intracellular accumulation, primarily due to increased efflux. Further, drug transport kinetics demonstrate the involvement of efflux mediated resistance, which was found to be reversed in the presence of efflux pump inhibitors (EPIs). Gene transcript analysis suggests differential upregulation of multiple efflux pumps across the genome of the mutant. Overexpression of one of the upregulated efflux pumps *Msmeg*_5659-5661, partially explains the XDR phenotype of the mutant, while also suggesting that the contribution of other efflux pumps is significant. Whole-genome sequencing (WGS) of Nor^r^ reveals that a mutation in *sox*R, a transcriptional regulator, could be responsible for the upregulation of the *Msmeg*_5659-5661 efflux pump by direct regulation, and other efflux pumps via indirect regulation. Thus, the present work demonstrates that high resistance to multiple drugs can arise even when the *Mycobacterium* was subjected to a single selection pressure. Further, alterations in drug transport is an important mechanism that leads to resistance to multiple drugs simultaneously.

## INTRODUCTION

The emergence of drug-resistant *Mycobacterium tuberculosis* (*M. tuberculosis*) poses a significant threat to tuberculosis treatment. Global tuberculosis report (2019) by WHO reported that in the year 2018, 51% world population with bacteriologically confirmed TB also tested positive for drug resistance. While about 46% of these were new cases, 83% of them were previously treated TB cases. Increase in multidrug-resistant TB (MDR-TB) cases has also been observed, i.e., resistance to two of the first-line agents used in TB treatment (isoniazid and rifampicin).^1^ On an average, about 9% of patients with MDR-TB had extensively drug-resistant TB (XDR-TB), where in addition to resistance against first-line anti-TB drugs, the bacteria is also resistant to fluoroquinolones and at least one of the three injectable second-line drugs (i.e., amikacin, kanamycin, or capreomycin).

Resistance to anti-TB drugs is mainly believed to be mediated by chromosomal mutations in target genes or bacterial enzymes that activate the pro-drugs.^2^ Since 1990, various mutations have been reported in clinical isolates that are resistant to anti-TB drugs.^2^ Rifampicin (RIF), a first-line anti-TB agent, inhibits bacterial RNA synthesis. Mutations that results in resistance to rifampicin are within the β subunit of RNA polymerase, encoded by the *rpo*B gene. Most of the mutations responsible for RIF resistance are within the 81-bp region of *rpo*B, referred to as the rifampicin resistance determinant region (RRDR).^3^ Another first-line anti-TB drug, isoniazid (INH) is a pro-drug, which on activation by *kat*G, a catalase and peroxidase enzyme, results in blocking of cell wall synthesis.^4^ Mutations associated with isoniazid resistance majorly affect *kat*G gene at codon 315 and the promoter region of *inh*A gene.^5^ Fluoroquinolones (FQs) are second-line drugs utilized for the treatment of drug-resistant TB or when the patient is intolerant to one of the first-line drugs.^6^ The FQs class of drugs inhibits nucleic acid synthesis by inhibiting the DNA gyrase enzyme encoded by *gyr*A and *gyr*B genes. Mutations in quinolone resistance determining region (QRDR) of *gyr*A and *gyr*B genes lead to FQ.^7^

Interestingly, target mutations do not explain all the cases of drug-resistant clinical isolates. For example, around 5 % of RIF-resistant *M. tuberculosis* clinical isolates lack mutations in the RRDR region of *rpo*B gene.^2, 8^ Similarly, 20-30% of INH-resistant *M. tuberculosis* clinical isolates do not harbor mutations in any of the known drug target genes associated with INH resistance. In contrast, up to 58% of FQ-resistant *M. tuberculosis,* clinical isolates do not have known resistance mutations.^2-3^ In a recent study, a sizeable fraction of ofloxacin-resistant clinical strains from Taiwan did not contain any mutations in either the *gyr*A or *gyr*B gene.^9^ Development of multiple drug resistance in *Mycobacterium* suggests that in addition to target mutations, additional mechanisms of drug resistance in mycobacteria also contribute to a substantially high level of resistance.^2^

Among the various drugs used for the treatment of tuberculosis, the fluoroquinolones class of drugs has the highest incidence of resistant strains where target gene mutations are absent. Resistance due to mechanisms other than target gene alteration also opens the possibility of multidrug resistance. In this study, we have characterized an in-vitro created norfloxacin resistant strain of *M. smegmatis* that contained no mutations in the *gyr*A and *gyr*B genes. We have investigated whether exposure to a single drug as a selection pressure can result in cross-resistance to other drugs. Using a combination of biochemical assays that shed light on the transport kinetics of drugs into/out of the cells, whole genome sequencing and expression studies, we determine the genomic basis of resistance. Understanding the mechanisms leading to multiple drug resistance will help us to develop new therapeutic agents against drug-resistant *Mycobacterium*.

## RESULTS AND DISCUSSION

### Preliminary characterization of a laboratory-selected mutant of *M. smegmatis* shows resistance to multiple drugs

Exposure to inadequate or sub-optimal drug dosage often results in the emergence of acquired drug resistance. To mimic this clinical situation, an FQ resistant mutant of *M. smegmatis* was generated using norfloxacin as the selection pressure. Wild-type *M. smegmatis* mc^2^ 155 strain (WT) was serially exposed to increasing concentrations of norfloxacin, ranging from a sub-inhibitory concentration of 0.5 µg/ml to concentrations up to 16 µg/ml (4x Minimum inhibitory concentration (MIC)). Subsequently, a norfloxacin-resistant mutant (Nor^r^ mutant) was isolated for further characterization. The MIC of norfloxacin against the Nor^r^ mutant was found to be 32 µg/ml, an increase of 16-fold as compared to the wild type To assess the cross-resistance potential of the Nor^r^ mutant, we determined its susceptibility to various other fluoroquinolones, including ofloxacin, ciprofloxacin, and moxifloxacin. The Nor^r^ mutant was found to be resistant to all the tested fluoroquinolones compared to the WT (Table 1), with maximum resistance towards norfloxacin.

**Table 1:**
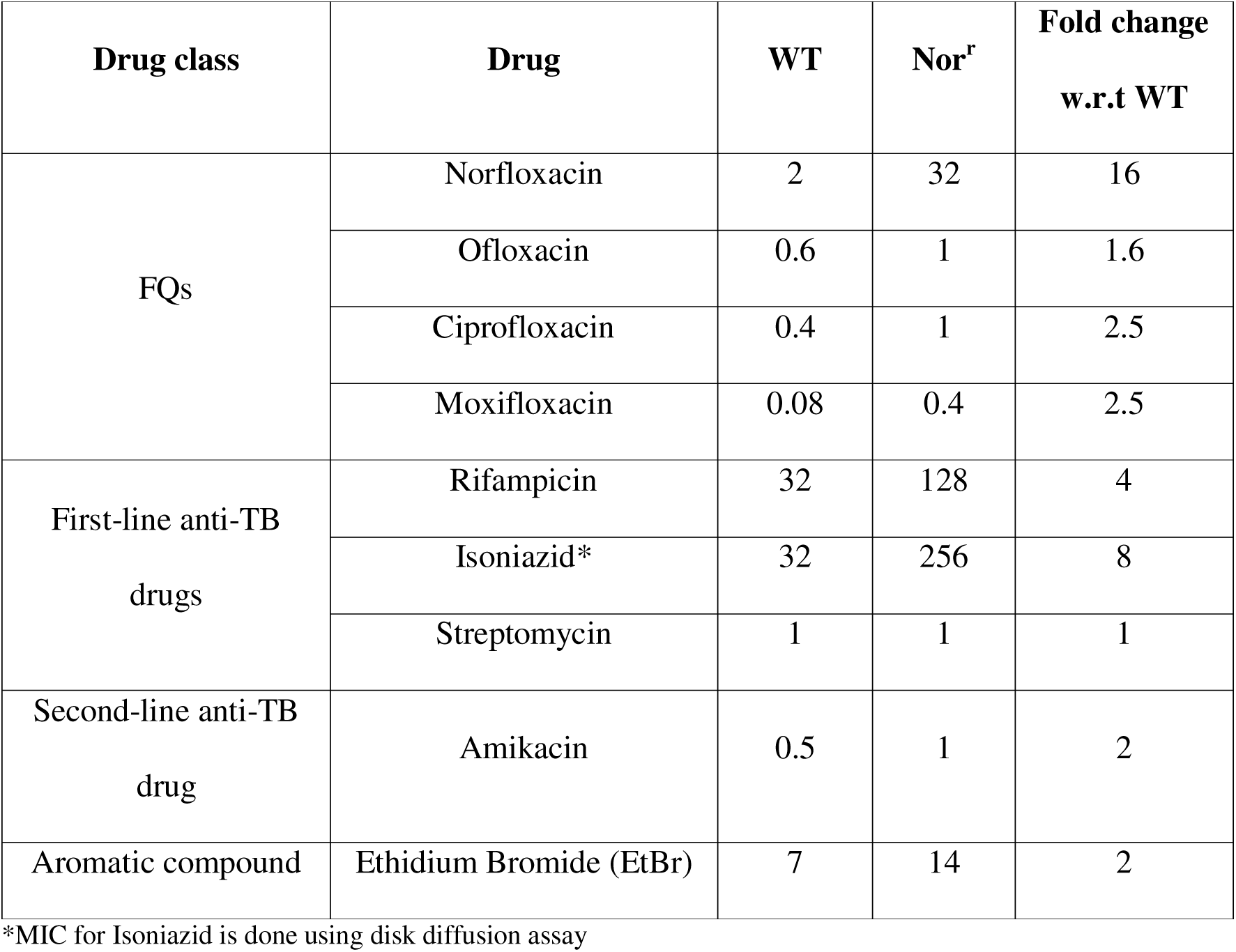
MIC of WT and Nor^r^ mutant of *M. smegmatis* to various fluoroquinolones (FQs), first-line, and second-line drugs.

When screened for resistance against other anti-TB drugs, Nor^r^ mutant was found to be 4-8-fold resistant to the first-line drugs, rifampicin and isoniazid, and 2-fold resistant to a second-line drug, amikacin (Table 1). Thus, the Nor^r^ mutant can be considered as an extensively drug-resistant (XDR) mutant of *M. smegmatis*.

To evaluate if the resistance phenotype led to any fitness disadvantage, growth kinetics of the Nor^r^ mutant and WT was studied. In the absence of norfloxacin, WT, and Nor^r^ mutant exhibited a similar doubling time (Figure 1).

**Figure 1:**
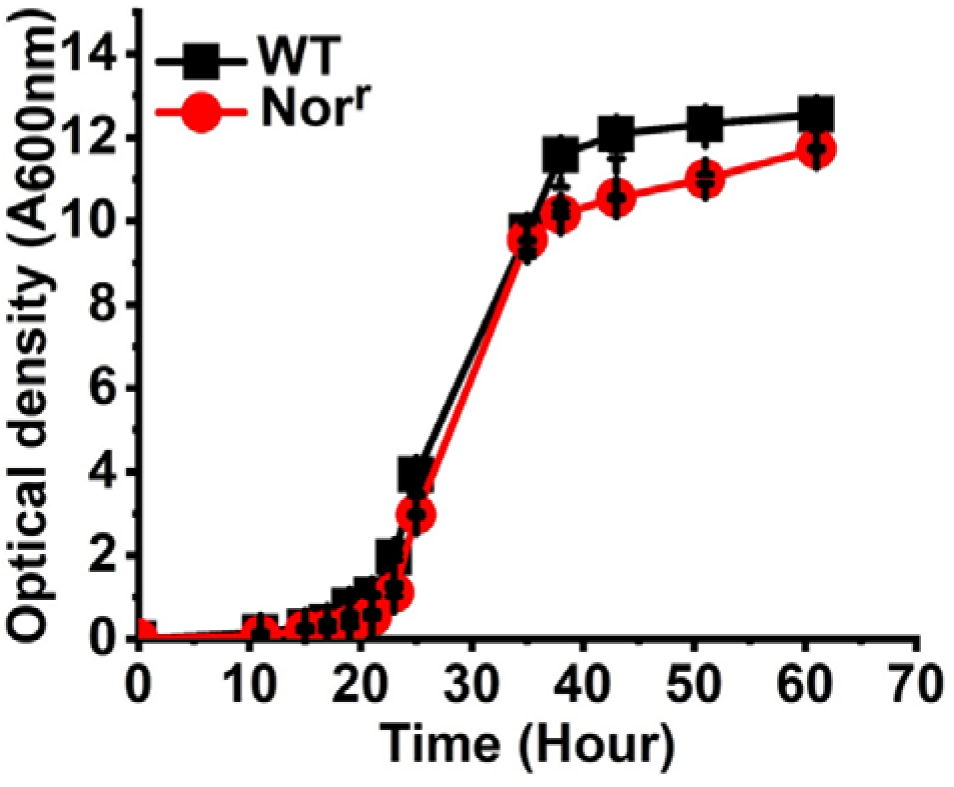
Comparison of WT and Nor^r^ mutant growth profile in the absence of the drug. (Note: O.D. values in the graph are the actual values in the culture medium, which were diluted below O.D.-1.0 for measurement.)

### Absence of mutations in any target genes

One of the primary mechanisms responsible for multiple-drug resistance in *M. tuberculosis* is the accumulation of multiple target mutations, each of which corresponds to the individual drug target genes to which the mycobacteria are resistant.^10^ The primary mechanism of FQ resistance is mutations in DNA gyrase. Numerous mutations have been reported in the quinolone resistance-determining region (QRDR) of gyrase A (*gyr*A) and gyrase B (*gyr*B) subunits of DNA gyrase associated with fluoroquinolone resistance.^11^ Therefore, we sequenced selected genes from the Nor^r^ mutant and compared it with that of the wild type strain. Surprisingly, no mutations were detected in the QRDR regions of either *gyr*A or *gyr*B genes from the Nor^r^ mutant. Although the mutant was resistant to rifampicin as well as isoniazid, no mutations were found in either of the respective target genes, *rpo*B, and *kat*G, in the mutant.

### Reduced intracellular drug levels in Nor^r^

The absence of target mutations clearly shows that other mechanisms of resistance are involved in conferring resistance to the Nor^r^ mutant against multiple drugs. Figure 2a shows the intracellular levels of norfloxacin at the end of one hour, in both Nor^r^ mutant and WT. The intracellular levels of norfloxacin were always lower by 1.5-3 folds in the Nor^r^ mutant as compared to WT for varying extracellular norfloxacin concentrations. At 2 μ (MIC of norfloxacin in WT), the intracellular norfloxacin amount was measured to be 100 ng/mg dry cell weight. Similar intracellular levels are achieved in the Nor^r^ mutant only at extracellular concentrations of more than 8 μg/ml. Since the Nor mutant was also resistant to rifampicin, we measured the intracellular rifampicin concentration in the mutant. For an extracellular rifampicin concentration of 32 µg/ml, the uptake of rifampicin in Nor^r^ mutant was reduced by 2-fold in comparison to WT (Figure 2b). Interestingly, in addition to various anti-TB drugs, Nor^r^ mutant was also found to be resistant to EtBr by 2-folds. The MIC of EtBr was 14 μg/ml against Nor^r^ mutant in comparison to 7 μg/ml against WT. EtBr emits a weak fluorescence in an aqueous environment outside the bacterial cells and becomes strongly fluorescent when it enters inside cells.^12^ Figure 2c shows the accumulation of EtBr in the Nor^r^ mutant as compared to WT. Analogous to norfloxacin uptake, EtBr accumulation was also severely reduced in the Nor^r^ mutant in comparison to WT (Figure 2c).

**Figure 2:**
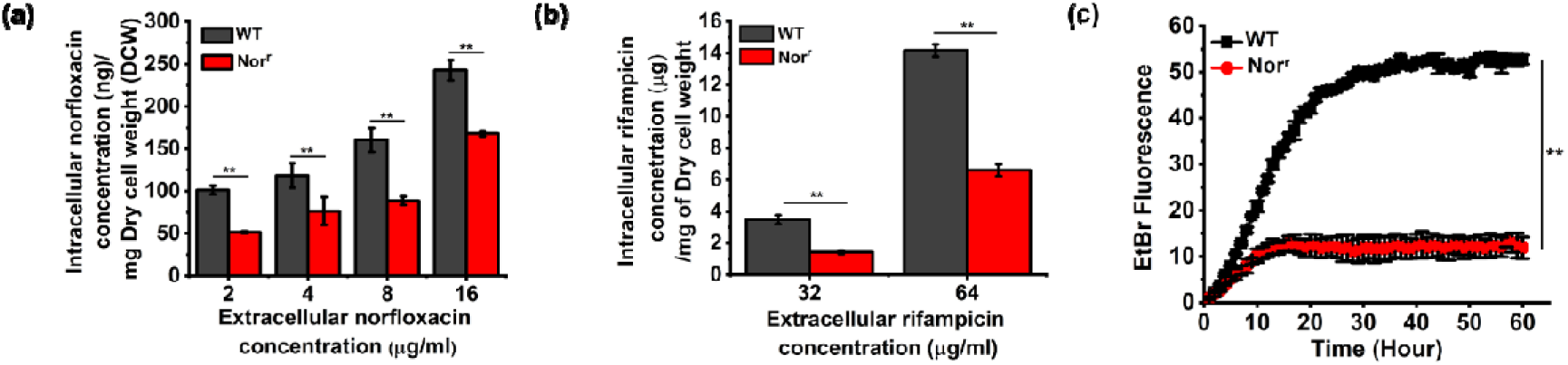
Comparison of the intracellular concentration of drugs in wild type and Nor^r^. Uptake of (a) norfloxacin and (b) rifampicin was compared for various extracellular concentrations of the corresponding drug. (c) EtBr (3µg/ml) accumulation in Nor^r^ mutant and WT. p-value < 0.001 denotes significance.

### Reversal of drug resistance in the presence of an efflux pump inhibitor suggests efflux as a mechanism of multiple drug resistance in Nor^r^

Low intracellular drug concentrations could be either due to lowered drug uptake and/or enhanced drug efflux. In order to investigate whether impaired drug influx is responsible for drug resistance in Nor^r^ mutant, we measured the membrane permeability of Nor^r^ mutant by NPN assay. As seen in Figure 3a, the influx of the fluorescent probe, NPN, is similar in the Nor^r^ mutant and WT, implying that membrane permeability is not altered in Nor^r^ mutant. Thus, the reason for decreased drug accumulation in Nor^r^ mutant could be enhanced efflux of drugs from the cells.

**Figure 3:**
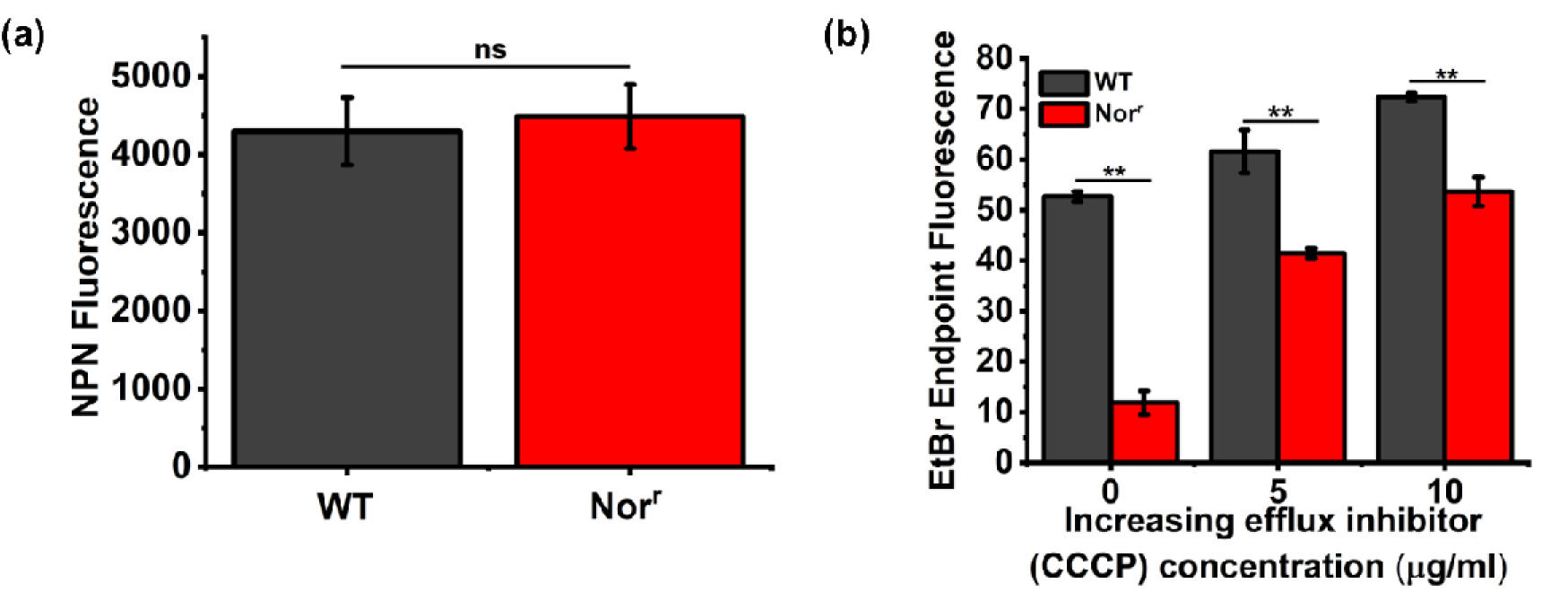
Reduced intracellular drug concentration in Nor^r^ mutant. (a) 1-N-phenylnaphthylamine (NPN) uptake as a measure of drug influx in Nor^r^ mutant and WT (b) EtBr accumulation in Nor^r^ mutant and WT in (b) presence of an EPI, CCCP. Accumulation of EtBr is dependent on both the uptake of the drug by diffusion and efflux by efflux pumps. ** represents p-value < 0.001, ns represents p-value >0.01

To further confirm the role of efflux in drug resistance of Nor^r^ mutant, EtBr accumulation in the mutant was measured in the presence of an efflux pump inhibitor (EPI), carbonyl cyanide m-chlorophenylhydrazone (CCCP). EtBr is a known efflux pump substrate and is hence used to mimic efflux of drugs in *M. smegmatis*.^13-15^ The accumulation of EtBr increased in the Nor^r^ mutant in the presence of increasing CCCP concentration (Figure 3b). At 10 µg/ml of CCCP, EtBr accumulation in Nor^r^ mutant increased up to the levels in WT. Correspondingly, the MIC of EtBr against the mutant reduced by four-fold in the presence of CCCP. Thus, enhanced uptake of EtBr by Nor^r^ mutant in the presence of CCCP suggests that active efflux is responsible for the extensive drug-resistant characteristic of the mutant.

Increased accumulation in the presence of CCCP led to a decrease in the MIC of the EtBr. To check whether increased efflux activity can explain the higher MIC of other drugs in Nor^r^, the MIC of various drugs, against the wild type and Nor^r^ mutant strain, was measured, in the presence of EPIs. In addition to CCCP, Sodium Orthovanadate (SOV) and Reserpine (RES) were also included (Table 2). CCCP is a protonophore which inhibits efflux activity by reducing the transmembrane potential and ATP production.^16^ CCCP affects pumps belonging to the major facilitator super family (MFS family) and to some extent the ATP binding cassette transporter family (ABC family).^17^ The EPI, SOV is a vanadium compound, which has the ability to block the flux of ions across the cell membrane by inhibition of Na^+^, K^+^-ATPase and Ca^2+^-ATPase thus leading to inhibition of ATPase and ATP efflux system. In addition, reserpine (RES), a plant-based EPI targets the resistance nodulation division (RND) superfamily and MFS family proteins.^16^ In the presence of EPIs, the MIC of most drugs decreased by 2-4 folds, both against the WT and the Nor^r^ strain, confirming our hypothesis that enhanced efflux activity is one the major reasons for the multidrug phenotype of Nor^r^.

**Table 2:** MIC of WT and Nor^r^ mutant of *M. smegmatis* to various fluoroquinolones (FQs), first-line and second-line drugs in the presence of EPIs (CCCP (10 µg/ml); SOV (20 µg/ml); RES (100 µg/ml)). The EPIs are chosen at 0.4x-0.5x MIC concentration.

| Drug class | Drug (µg)* | WT |  |  |  | Nor<sup>r</sup> |  |  |  |
| --- | --- | --- | --- | --- | --- | --- | --- | --- | --- |
|  |  | Without EPI | With EPI |  |  | Without EPI | With EPI |  |  |
|  |  |  | CCCP | SOV | RES |  | CCCP | SOV | RES |
| FQs | NOR | 2 | 1 | 0.5 | <0.25 | 32 | 8 | 16 | 16 |
|  | OFL | 0.6 | 0.3 | 0.3 | 0.3 | 1 | 0.5 | 0.25 | 0.625 |
|  | CIP | 0.4 | 0.2 | 0.1 | <0.05 | 1 | 0.5 | 0.5 | <0.625 |
|  | MOXI | 0.08 | 0.04 | 0.02 | 0.02 | 0.4 | 0.2 | <0.05 | <0.05 |
| First-line anti-TB drugs | RIF | 32 | 16 | <4 | <4 | 128 | 64 | 32 | <32 |
|  | INH* | 32 | 16 | 16 | 16 | 256 | 64 | 128 | 64 |
|  | STP | 1 | 0.5 | 0.5 | <0.625 | 1 | 0.5 | 0.5 | 0.25 |
| Second-line anti-TB drug | AMK | 0.5 | 0.25 | 0.25 | <0.625 | 1 | 0.25 | 0.5 | 0.25 |
| Aromatic compound | EtBr | 7 | 3.5 | 3.5 | <0.875 | 14 | 3.5 | 7 | 7 |

\*NOR-Norfloxacin, OFL-Ofloxacin, CIP-Ciprofloxacin, MOXI-Moxifloxacin, RIF- Rifampicin, INH-Isoniazid, STP-Streptomycin, AMK-Amikacin, EtBr-Ethidium bromide

Interestingly, the three EPIs had differential effects on the bacteria depending on the drug class. For most drugs tested, atleast one of the EPIs was able to reduce the MIC of the Nor^r^ mutant to WT levels. However, norfloxacin and isoniazid were exceptions. In the presence of CCCP, the MIC of norfloxacin against the mutant was 8 µg/ml, which is four-fold higher than that against the WT. Similarly, both CCCP and RES were able to reduce the MIC of isoniazid against the mutant to 64 µg/ml.

### Whole-genome Sequencing

The data so far suggests that the MDR phenotype of the Nor^r^ mutant can be explained partially by enhanced efflux in the mutant. To understand the molecular basis of this, whole-genome sequencing was performed. Table 3 lists the primary mutations found in the Nor^r^ mutant genome. One mutation that is important to note is that in the *sox*R transcriptional regulator. Mutations in this gene have been shown to lead to overexpression of efflux pumps in many other bacteria and thus lead to resistance to drugs.^18^ The targets of *sox*R in the *M. smegmatis* genome have not been delineated yet. However, a previous study had reported the presence of sox-box at three locations in the genome (http://soxrbox.mit.edu/); one of these is upstream of *sox*R. The other two are upstream of MSMEG_5659-61 genes that code for an ABC transporter and *sig*H4 (MSMEG_0574) gene that encodes for a sigma factor. Measurement of mRNA levels in the Nor^r^ mutant indeed revealed that both the genes are constitutively overexpressed in the Nor^r^ mutant with a log_2_ ratio of 2-fold for *Msmeg*_5659 and 10-fold for *Msmeg*_0574.

**Table 3.** Unique mutations* in Nor^r^ Mutant w.r.t WT population through Illumina platform.

| Position | Gene | Protein | Mutation type | Effect of mutation | SDP/DP |
| --- | --- | --- | --- | --- | --- |
| 922311 | MSMEG_0836 | Carboxylate-amine ligase | SNP | Synonymous | 156/143 |
| 2318499 | MSMEG_2237 | Anaerobic dehydrogenase | SNP | (V659A) | 145/143 |
| 5535979 | *soxR* (MSMEG_5450) | Redox-sensitive transcriptional activator | SNP (gGc-gAc) | (G113D) | 132/120 |
| 886311 | MSMEG_0862 | hypothetical protein | INDEL (Deletion of G) |  | 128/112 |

\*Frequency >50%

### Over-expression of *Msmeg*_5659-61 efflux pump can partially explain the extreme-drug resistance phenotype

To investigate the contribution of the *Msmeg*_5659-61 efflux pump to Nor^r^ resistance, we expressed an additional copy of the *Msmeg*_5659-61 genes in *M. smegmatis* mc^2^155 under a constitutive promoter, to create the strain *Msmeg*_5659_OE_. Table 4 presents the MIC of the various drugs against *Msmeg*_5659_OE_. The construct with the empty plasmid was used as a positive control. Over-expression of *Msmeg*_5659 in multiple copies leads to increased resistance of the strain to fluoroquinolones. In addition, the MIC of rifampicin and isoniazid also increased by 4-8 folds. Interestingly, the MIC of norfloxacin against *Msmeg*_5659_OE_ was 16 µg/ml as opposed to the observed MIC of 32 µg/ml against the Nor^r^ mutant. However, the mRNA expression level of MSMEG_5659 was higher in *Msmeg*_5659_OE_ than that observed in the Nor^r^ mutant (Figure 4b). Despite the higher transcript levels of *Msmeg*_5659 in the construct, the observed MIC of Norfloxacin against *Msmeg*_5659_OE_ is lower compared to that against the Nor^r^ Mutant.

**Figure 4:**
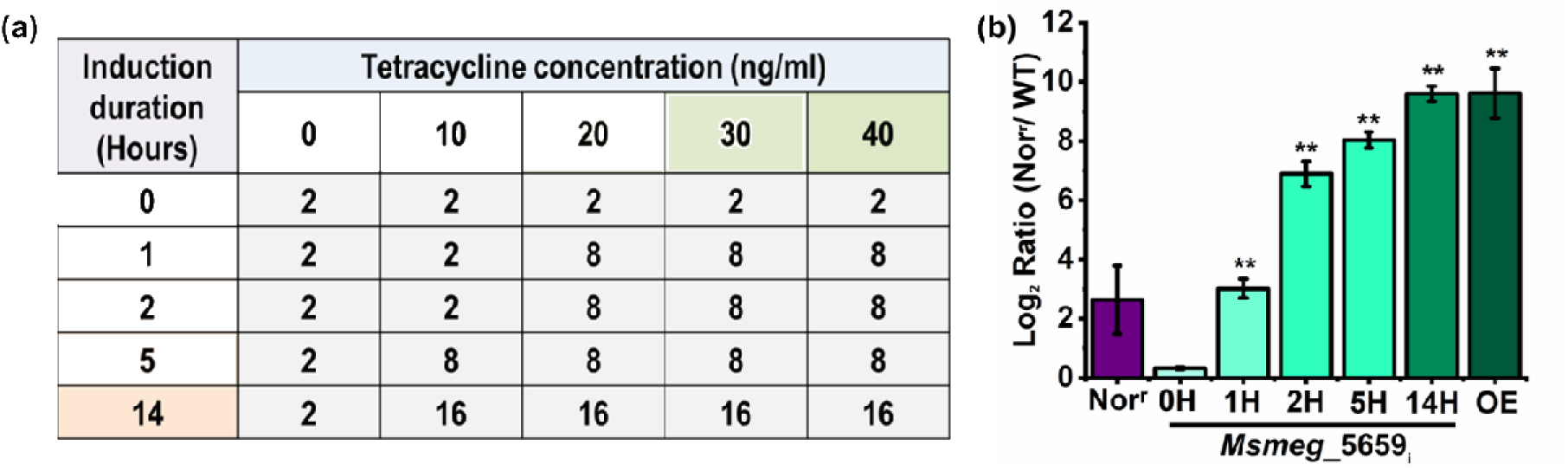
Effect of varying expression of MSMEG_5659 on MIC of norfloxacin. (a) MIC of Norfloxacin against *Msmeg*_5659_i_ under varying inducer concentrations and induction time. (b) Transcript levels of *Msmeg*_5659 in various strains with respect to the WT levels. The log2 ratios are shown for the Nor^r^ mutant, *Msmeg*_5659_i_ and Msmeg_5659_OE_ strains (OE). SigA was used as the housekeeping gene for normalization of qRT data. *Msmeg*_5659_i_ was induced with 40 ng/ml of tet for varying times (0 to 14 hours) as mentioned, and corresponding mRNA levels were measured. 0H refers to sampling just prior to induction; 1H refers to cells induced for 1-hour etc. p-value < 0.001 denotes significance.

**Table 4:** Effect of over-expression of efflux pump on drug susceptibility in construct *Msmeg*_5659_i_

| Drug class | Drug | WT/<br>EV<sub>OE</sub> | Nor<sup>r</sup><br>mutant | *Msmeg_5659*<sub>OE</sub> | *Msmeg_5659<sub>i</sub>*<br>(40ng/μl for<br>1 Hr) | *Msmeg_5659<sub>i</sub>*<br>(40ng/μl for<br>14Hrs) |
| --- | --- | --- | --- | --- | --- | --- |
|  |  | MIC (μg/ml) |  |  |  |  |
| FQs | Norfloxacin | 2 | 32 | 16 | 8 | 16 |
|  | Ofloxacin | 0.6 | 1 | 2.4 | 2.4 | 4.8 |
|  | Ciprofloxacin | 0.4 | 1 | 3.2 | 0.4-0.8 | 1.6 |
|  | Moxifloxacin | 0.08 | 0.4 | 0.16 | 0.16 | 0.16 |
| 1<sup>st</sup> line | Rifampicin | 32 | 128 | 128 | 64 | 512 |
|  | Isoniazid* | 32 | 256 | 256 | 32 | 256 |
|  | Streptomycin | 1 | 1 | 2 | 4-8 | 16 |
| 2<sup>nd</sup> line | Amikacin | 0.5 | 1 | 4 | 0.5-1 | 4 |
| Aromatic | EtBr | 7 | 14 | 14 | 14 | 14 |

To determine the effect of efflux pump transcript levels on the observed MIC, we cloned *Msmeg*_5659 under a tet-inducible promoter and named the construct as Msmeg_5659_i_. The concentration of inducer, tetracycline, and the induction time were varied to determine their effect on the MIC of norfloxacin (Figure 4a). Simultaneously, the mRNA levels of *Msmeg*_5659 were also measured (Figure 4b). Data for other drugs is presented in Table S3. The MIC increases as a function of both the time of induction as well as the tetracycline concentration. We observed a four-fold higher resistance at an inducer concentration of 20 ng/ml after an hour of induction. A similar four-fold higher resistance was observed with an inducer concentration of 10 ng/ml after five hours of induction. Inducing the cells for 14 hours with tetracycline led to the maximum observed MIC of 16 µg/ml. The transcript levels also increased with increasing induction time with maximum levels similar to that achieved in the *Msmeg*_5659_OE_ cells. Thus, over-expression of this efflux pump leads to increased resistance of the bacteria to a diverse range of drugs including the FQs, first-line and second-line drugs. However, knocking out *Msmeg*_5659 does not affect the susceptibility of cells towards any of the drugs tested (Table S4). This could be attributed to the large number of efflux pumps that are present in the *M. smegmatis* genome and extensive cross-talk between them.

### Additional efflux pumps are upregulated in drug-resistant Nor^r^

Overexpression of MSMEG_5659 could only partially explain the resistance profile of Nor^r^ mutant. *M. smegmatis* has a large number of transporters (greater than 400), out of which more than 120 are predicted to be responsible for multidrug efflux.^2^ However, very few of them have been characterized so far. LfrA, an efflux pump from the major facilitator superfamily (MFS), is responsible for the efflux of FQs.^19^ It is also known to be involved in EtBr and rifampicin resistance. Thus, we quantified the mRNA expression level of *lfr*A in the WT and Nor^r^ mutant (Figure 5). Besides, expression levels of additional efflux pumps were also measured, selected based on prior literature or their homology to known *M. tuberculosis* efflux pumps.

**Figure 5:**
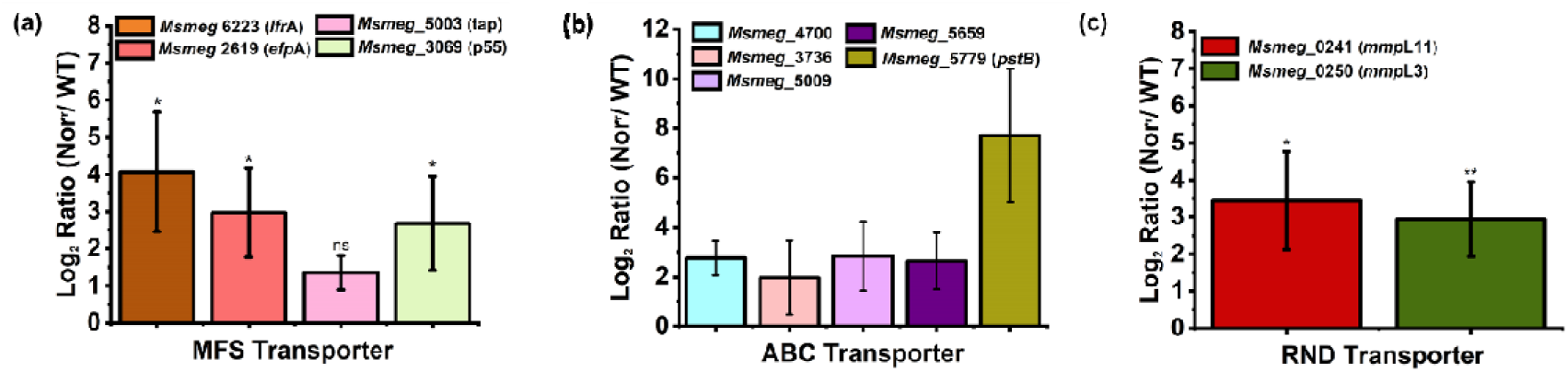
mRNA expression level of various classes of transporters in the Nor^r^ mutant wrt wild type. (a) MFS transporters (*lfr*A, *efp*A (Rv2846c)), *tap* (Rv1258c)), and *Msmeg*_3069 (p55(Rv1410c)), **(b)** ABC transporters Msmeg_5659 (Rv0194), *Msmeg*_5779 (*pst*B (Rv0933)), *Msmeg*_5009 (Rv1272c), *Msmeg*_3763 (Rv1686c), and *Msmeg*_4700 (Rv2477c) (b) RND transporters (*mmp*L 11, and 3). The corresponding homolog in *M*. *tuberculosis* H37Rv is listed in bracket. * represents p < 0.05, ** represents p-value < 0.001, ns represents p-value >0.01

lfrA was found to be constitutively overexpressed in the Nor^r^ mutant with a log_2_ ratio of 4 with respect to the the wild type. *Msmeg*_2619 and *Msmeg*_3069 (homologous of *efp*A and *p*55 efflux pumps from the MFS family) are also upregulated by more than 2 fold (log_2_ ratio >1) in the mutant. *efp*A, and *p*55 efflux pump genes are responsible for resistance against isoniazid, streptomycin, and tetracycline in *M. tuberculosis*^20-22^. Many efflux pumps belonging to the ATP-binding cassette superfamily (ABC) transporter family, such as *pst*B, *Msmeg*_5659, *Msmeg*_4700, *Msmeg*_5009, *Msmeg*_5779, and *Msmeg*_3736 and two pumps (mmpL11 and mmpL3) from the RND superfamily were also constitutively over-expressed by more than 2 fold (log_2_ ratio >1) in the Nor^r^ mutant. *pst*B, is involved in the efflux of isoniazid and rifampicin in *M. tuberculosis* and *M. smegmatis.*^23^ Efflux pump mmpL3 is involved in isoniazid resistance in *M. tuberculosis*.^21, 24^ Disruption of mmpL11 in *M.tb* leads to attenuation of growth in mice lungs^25^ and delayed heme uptake.^26^ mmpL3 and mmpL11 are found to be conserved across all mycobacterial genome sequenced so far, suggesting a fundamental role in mycobacterial biology.^27^ Tap and *Msmeg*_5659 have been found to be upregulated in the presence of Streptomycin.^28-30^ and over expression of Tap leads to 4 fold increased resistance of Streptomycin in *Mycobacterium bovis* BCG.^29^ Homologs of most of these efflux pumps have been shown to be upregulated in the Beijing clinical isolates with a fold change range of 3-10.^31^ Thus, the multidrug-resistance characteristic of Nor^r^ mutant can be attributed to the reduced uptake of drugs due to the constitutive upregulation of these efflux pumps.

### Nor^r^ mutant is also resistant to oxidative stress

Mutations in the *Sox*R protein have been shown to lead to efflux pump over-expression. In addition, *sox*R affects the survival of the bacteria in the presence of redox-cycling drugs.^32^ We therefore quantified the MIC of a few redox-changing compounds against both WT and Nor^r^. Increased resistance of Nor^r^ mutant to plumbagin and hydrogen peroxide (H_2_O_2_) as compared to that of the WT, suggests that the Nor^r^ mutant possesses improved defense towards oxidative stress as compared to that of WT (Table 5). The resistance of Nor^r^ to norfloxacin could not be completely reversed using CCCP or another EPI. We hypothesize that the observed resistance of Nor^r^ towards norfloxacin could be due to a combination of increased efflux activity as well as increased quenching of reactive oxygen species. In order to further establish the role of Reactive oxygen species (ROS) quenching in the mutant, we used an antioxidant in the presence of a ROS producing FQ. If the Nor^r^ mutant has a higher ROS quenching ability due to mutations in the *sox*R gene, the inhibitory concentration of a ROS producing drug will increase in the presence of antioxidants.^33^ In the presence of ascorbic acid, the norfloxacin MIC against WT increased by 4-fold to 8 μg/ml. Similarly, in the presence of ascorbic acid, the MIC of norfloxacin against Nor^r^ also increases further. Note that individual MIC for ascorbic is 1.6 mg/ml (Table 5). This significant increase in the MIC strongly suggests the involvement of ROS along with the upregulated efflux pumps leads to increased resistance in the Nor^r^ mutant.

**Table 5:** MIC of ROS producing drugs/ compounds against WT and Nor^r^ mutant.

| Strain | Individual inhibitory concentrations |  |  |  | Inhibitory concentrations in combination |  |
| --- | --- | --- | --- | --- | --- | --- |
|  | Plumbagin | Hydrogen peroxide (H<sub>2</sub>O<sub>2</sub>) | Norfloxacin | Ascorbic acid | Norfloxacin | Ascorbic acid |
| WT | 2 | 2.5 | 2 | 1600 | 8 | 200 |
| Nor<sup>r</sup> | 16 | 5 | 32 | 1600 | 64-128 | 200 |

Other than norfloxacin, the MIC of isoniazid could not be reversed to levels similar to WT in combination with any of the three EPIs tested. This suggests that increased resistance of Nor^r^ towards isoniazid could also be due to a combination of efflux pump over expression and reduced production of reactive species. Both the strains produce similar ROS levels in the absence of any drug. However, the Nor^r^ mutant produces much lower ROS as compared to the wild type when treated with isoniazid (Figure 6). The reduced ROS production could be a combination of the reduced intracellular accumulation of INH as well as the increased ability of Nor^r^ mutant to quench the ROS.

**Figure 6:**
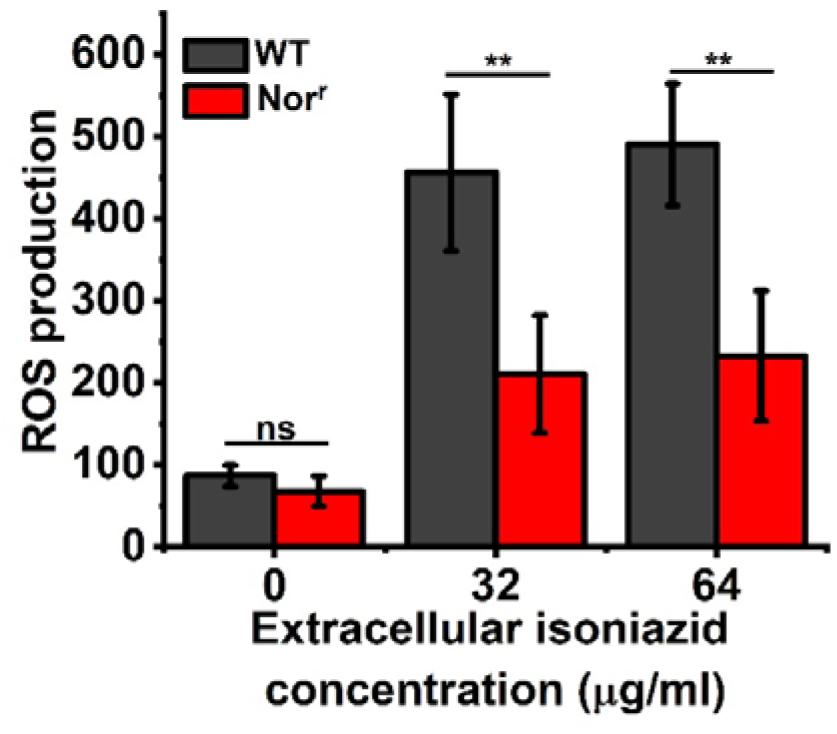
Comparision of production of ROS in the WT and Nor^r^ on treatment with isoniazid at various concentrations. ** represents p-value < 0.001, ns represents p-value >0.01.

## Discussion

XDR-TB is of grave concern, especially with patients that are HIV positive. There is a critical need to understand the evolution and spread of resistance to multiple antibiotics and thereby devise solutions. In this study, an in-vitro laboratory mutant of *M.smegmatis* has been characterized that exhibited characteristics similar to XDR-TB.

The norfloxacin-resistant mutant (Nor^r^ mutant) was not only resistant to norfloxacin but also resistant to a panel of other fluoroquinolones. Interestingly, while the mutant was created with norfloxacin alone as a selection pressure, the resultant mutant was resistant to first-line drugs, rifampicin and isoniazid, and second-line drug amikacin, thus displaying characteristics of XDR-TB. Further, in the absence of any drug pressure, the mutant growth kinetics was similar to that of the wild-type. Fitness costs play an essential role in the spread of multi-drug resistance.^34, 35^ Similar growth kinetics suggests that the increased resistance of the evolved strain does not impose any fitness cost on it, and therefore should be able to survive with equal efficiency in the presence of the wild type strain. Growth rates of MDR-TB and XDR-TB strains of *M. tuberculosis* containing drug-resistant mutations have been reported to be similar to that of the susceptible strain.^36^

The primary mechanism of multiple drug resistance in *Mycobacterium* is described as mutations in the individual drug target genes. Hence, most studies characterizing MDR/XDR-TB from various geographic regions deal with target mutations.^37-42^ Accumulation of mutations in multiple genes, including *gyr*A, *gyr*B, *rpo*B, *kat*G, etc. have been reported in MDR/XDR strains. Recently many unique strains have been reported with no mutation in the typical hot-spots for various drugs, including RIF, INH, fluoroquinolones, and aminoglycoside.^38, 39^ Indeed, an alternate mechanism of resistance is now emerging. For example, the resistance of ofloxacin in isolates without any *gyr*A/B mutation was mapped to a novel mutation in the *ecc*C5 gene.^43^ In another study, multidrug resistance in *M. smegmatis* evolved under ciprofloxacin pressure was mapped to mutations in ribosomal genes.^44, 45^ Loss of the sigma factor, *sig*I, led to resistance of *M. tuberculosis* to isoniazid, without any fitness defects.^43^

Mutations in none of the target genes was detected in Nor^r^. Simultaneouly, the reduced intracellular drug concentration of norfloxacin, rifampicin and etbr, coupled with no change in membrane permeability suggested enhanced efflux activity to be responsible for resistance. This was confirmed by the decrease in resistance of Nor^r^ in the presence of EPIs. While MIC of most drugs against the Nor^r^ mutant could be restored to wild-type levels by the addition of EPIs, norfloxacin was an exception. The action of fluorquinolones is both due to inhibition of gyrA as well as due to generation of oxidative stress. The Nor^r^ mutant was found to be resistant to plumbagin and hydrogen peroxide. Therfore, the multidrug resistant characteristic of this strain is a combination of both enhanced efflux and increased resistance to oxidtaive stress.

To identify the genomic basis of this enhanced efflux activity and oxidtaive stress resistance, WGS of Nor^r^ was performed. One of the major mutations observed through WGS analysis of the Nor^r^ mutant was in the *sox*R gene, where a SNP (gGc-gAc) would lead to the glycine (G) at residue 113 being changed to aspartic acid (D). The G113D substitution, which alters a residue in the [2Fe-2S] cluster region, could render *sox*R in Nor^r^ mutant constitutively active.^46, 47^ While the role of *sox*R is not yet fully described in *M. smegmatis*, it’s role as an oxidative stress regulator has been demonstrated in many other organisms. In *E.coli*, *sox*R is a regulator of the *sox*S gene that in turns affects activity of many oxditiave stress response genes and also impacts multidrug resistance.^48, 49^ One such example is seen in clinical isolates of *E.coli* where a mutation in the *sox*R gene leads to multidrug resistance phenotype.^18^ SoxR has also been shown to directly or indirectly regulate efflux pumps in many organisms. In *Klebsiella pneumoniae*, mutation in *sox*R leads to over expression of *sox*S and the acrB efflux pump and corresponding increase in MIC of various drugs.^50^ The *sox*S gene is missing in many gram-positive bacteria, and *sox*R directly controls a small subset of genes. For example, in *P. aeruginosa*, it was observed that soxR directly activates a six-gene regulon, which includes efflux pumps.^51^ Similarly, the *sox*R gene of *Streptomyces coelicolor* also activates a six-gene regulon including an efflux pump.^52^

The increased efflux activity of Nor^r^ is due to constitutive upregulation of multiple efflux pumps. The *Msmeg*_5659-61 operon, encoding an ABC efflux pump is predicted to be a direct target of soxR, based on the presence of the soxBox, the soxR binding site. We hypothesize that the constitutive over-expression of MSMEG_5659-61 efflux pump in the Nor^r^ mutant is due to the constitutive activity of soxR. Expressing an additional copy of the *Msmeg*_5659-61 genes, clearly shows that the upregulation of this efflux pump can explain the increased resistance of Nor^r^ to different drugs. Interestingly, over-expression of the MSMEG_5659-61 efflux pump alone could not explain the resistance to most drugs, as much higher mRNA levels would be required to attain the MIC exhibited by the Nor^r^ strain. On measuring the transcript levels of additional efflux pumps, many of these were also constitutively over-expressed in Nor^r^. Absence of soxbox upstream of these genes suggested indirect measures of control. In addition to MSMEG_5659-61, the soxbox is also located upstream of the sigma factor, *sig*H4 (*Msmeg*_0574). We hypothesis that the sigma factor *sig*H4 is involved in the upregulation of multiple efflux pumps, further leading to the MDR phenotype in Nor^r^ Mutant. Further experiments are required to delineate the precise role of this gene in *M. smegmatis*. However, its upregulation in the presence of 7mM of H_2_O_2_ suggests that this sigma factor plays a role in the regulation of oxidative stress response.^20^ Homologues of this gene in M. *t*b have been shown to be essential for virulence.^20^ Genomic analysis through microarray suggests that *sig*H gene mediates the transcription of 30 genes directly and 150 genes indirectly in M. *tb*.^53^ including many transcriptional regulators, oxidative stress genes and an efflux pump. Future work will focus on understanding the role of soxR and sigH4 in *M. smegmatis* and their contribution to multidrug resistance.

Evolution under norfloxacin as selection pressure led to resistance to many drugs that are not structurally related. Results suggest that there is significant cross-talk among the efflux pumps and their regulator for the various anti-Tb drugs currently in use. This work assumes siginificance as FQs are essential drugs for tuberculosis therapy, and clinical trails are underway to design alternate regimes, including FQs that can reduce the treatment time. Thus, the evolution of resistance in Mycobacteria in the presence of FQs should be investigated throughly as this can impact the treatment regime.

## CONCLUSION

To summarise, this is the first study, which characterizes a laboratory mutant, possessing an extensive-drug resistant characteristic and proves that efflux is an essential mechanism of drug resistance. Through drug transport and gene expression studies, we have established that mechanisms other than target gene mutations can also confer an extensive-drug resistant phenotype to a laboratory-derived mutant of *M. smegmatis*. A schematic representation of various mechanisms of drug resistance in Nor^r^ mutant is elucidated in Figure 7. Thus, characterizing clinical XDR-TB isolates for such alterations could shed light on multidrug-resistant strains, which cannot be explained by classical target mutations. This study highlights the role of efflux as an essential mechanism for acquiring multiple drug-resistant phenotypes by mycobacteria.

**Figure 7:**
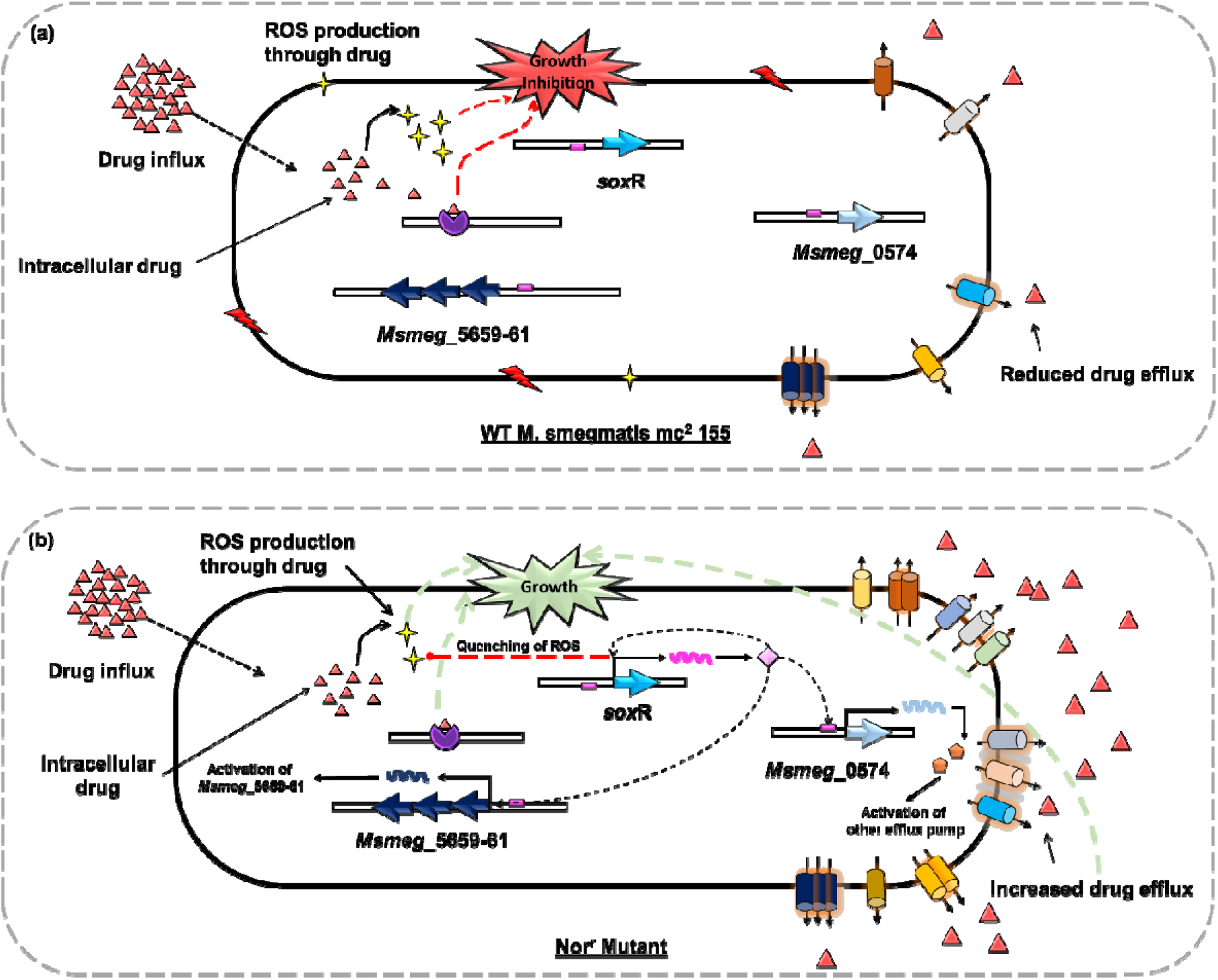
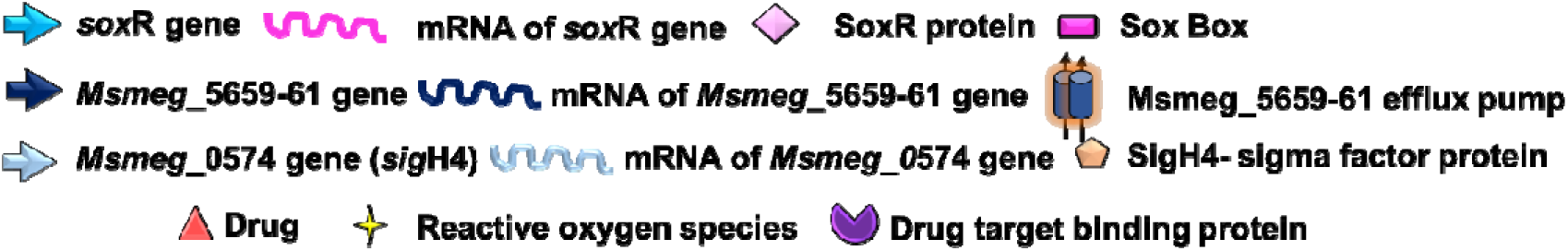
Proposed mechanism of extreme drug resistance of Nor^r^ mutant in comparison to WT *M. smegmatis*. (a) In the case of WT *M. smegmatis* strain, the inhibitory effect of the drug is due to a binding of drug to its target protein. For some drugs such as FQs, in addition production of ROS also leads to growth inhibition and cell death. (b) In the case of Nor^r^ Mutant, we propose that the strain utilizes multiple resistance mechanisms in order to survive. Intracellular influx of drug is similar to that of WT. However, upregulation of multiple efflux pumps leads to a decrease in the effective intracellular concentration of the drug. We propose that the mutation in soxR gene leads to its constitutive activity. While the *Msmeg*_5659-61 ABC transporter is directly regulated by soxR, other efflux pumps are potentially indirectly regulated via the sigH4 protein that is regulated by soxR. Presence of mutated soxR also leads to increased resistance to oxidative stress by quenching of the ROS. Together, this leads to increased survival of the Nor^r^ Mutant.

## MATERIAL AND METHODS

### Culture conditions

*Mycobacterium smegmatis* mc^2^155, *M*. *smegmatis* mc^2^ 155 harboring pSTK (overexpressing plasmid), and pSTKiT (Integrating-inducible plasmid) (pSTK (addgene plasmid #44560) and pSTKiT (addgene plasmid #44562) was a gift from vinay nandicoori)(Parikh, 2013 #89) used for the study was cultured in Middlebrook 7H9 (M7H9, Hi-media) media supplemented with 10% (v/v) ADC (5 g albumin, 2 g glucose and 0.85 g NaCl in 100ml distilled water) at 37°C and 180 rpm. ADC was initially filter-sterilized before addition to the autoclaved media. Luria-Bertani (LB) media (Hi-media) was used for growing the bacteria on an agar plate.

### Minimum inhibitory concentration (MIC)

MIC of rifampicin (RIF), isoniazid (INH), norfloxacin (NOR), ofloxacin, amikacin, streptomycin, ciprofloxacin, ethidium bromide (EtBr), carbonyl cyanide m-chlorophenylhydrazone (CCCP), Sodium orthovanadate (SOV), and Reserpine (RES) was determined by broth microdilution method, according to the clinical and laboratory standards institute (CLSI) guidelines.^54^, MIC of isoniazid was determined zone of inhibition assay. EPI for MIC was observed as 20 µg/ml for CCCP, 256 µg/ml for RES, and 100 µg/ml for SOV against both WT and Nor^r^ mutant. Kanamycin (selection pressure) and tetracycline (inducer) was present in the culture as well as dilution media while doing MIC experiments for *Msmeg*_5659_OE_ and *Msmeg*_5659_i_ respectively.

### Selection of norfloxacin resistant mutant (Nor^r^) of *M. smegmatis*

Mutant was generated by serially exposing the wild-type (WT) bacteria to agar media containing norfloxacin at a sub-inhibitory concentration of 0.5 µg/ml. Sub-optimal dose exposed bacteria were then exposed to increasing norfloxacin concentration, ranging from sub-inhibitory concentration (0.5 µg/ml and 1 µg/ml) to inhibitory concentration (2 µg/ml) and subsequently passaging on norfloxacin concentrations up to 16 µg/ml (higher than MIC). A single colony was selected from an agar plate containing 16 µg/ml norfloxacin and was subsequently cultured in liquid M7H9 medium supplemented with ADC (without any norfloxacin). This liquid culture was stored at -80°C as stock for further experiments.

Furthermore, to check the stability of the mutant in the absence of norfloxacin, Nor^r^ mutant from stock culture was further passaged in plain LB agar media (without containing norfloxacin) for ten passages. Subsequently, WT and mutant cells, from the 10^th^ passage, were streaked on an agar plate containing 8 µg/ml of norfloxacin. The Nor^r^ mutant was able to grow on norfloxacin containing plate even after ten passages on media without norfloxacin, whereas the WT failed to grow. Thus, the mutant was genetically stable and resistant to norfloxacin, even in the absence of antibiotic pressure.

### Intracellular rifampicin measurements

Intracellular rifampicin concentration was measured using the reverse phase –HPLC. 0.5 O.D. cells were concentrated to O.D. 5 and resuspended in Middlebrook 7H9 broth with desired concentrations of rifampicin. Treated cells were incubated for an hour at 37°C with 200 rpm shaking. Extracellular rifampicin was estimated to account for the intracellular concentration of rifampicin in the media after 1 hour. For analysis, samples were collected at 0 hours and 1 hour and centrifuged at 8500 rpm for 10 min. The supernatant was filtered through a 0.2-micron filter, and the pellet was dried for Dry Cell Weight (DCW) measurement. Samples were then passed through a 5 cm C18 column (Agilent make) in High-Performance Liquid Chromatography (HPLC) Agilent 1260 infinity system. A 50:50 mixture of water acidified to pH 2.27 with ortho-phosphoric acid and acetonitrile was used as the mobile phase. The flow rate used was 1.2 ml/min and elute measured with Ultraviolet (UV) detector at 333 nm.

### Estimation of ROS

The generation of reactive oxygen species (ROS) was measured using the dye 2’,7’-dichloroflurescein-diacetate (H_2_DCFDA) ^55^. In the presence of ROS, non-fluorescent dye, 2’,7’-dichloroflurescein-diacetate (H_2_DCFDA) is converted to highly fluorescent compound 2’,7’-dichloroflurescein (DCF) inside the cells. Briefly, 0.5 OD_600nm_ *M. smegmatis* WT and Nor^r^ mutant cells were incubated for an hour with the desired concentration of isoniazid. After an hour of incubation, H_2_DCFDA (final concentration as 10µM) was added to the cells for 10min. Subsequently, the cells were washed and resuspended in PBS and 10µM H_2_DCFDA. 200µl of this suspension was transferred to 96 well plates (black polysorp, Nunc). ROS generation was measured by measuring the fluorescence in a microplate reader (spectramax multimode M5, molecular devices) at 37°C using 488 nm and 525 nm excitation and emission wavelengths, respectively.

### Intracellular fluoroquinolone (Norfloxacin) measurements

Intracellular norfloxacin was quantified using a fluorescence-based method. Briefly, *M. smegmatis* WT and Nor^r^ mutant cells were grown till they reach OD_600nm_ of 0.5. They were further concentrated to 5 OD cells. To these cells, norfloxacin was added at different concentrations and incubated for 1 hour at 37°C at 180 rpm. After 1 hour of incubation, the cells were centrifuged at 10000 rpm at 4°C for 10 min and were washed twice with chilled PBS. Finally, the pellet was lysed by adding 0.1 M glycine HCl (pH 3) and kept overnight at room temperature. Samples were centrifuged, and supernatant fluorescence was measured using 281 nm and 440 nm as excitation and emission wavelengths, respectively. A standard fluorescence curve of varying norfloxacin concentrations in glycine HCl was plotted. Results were expressed in ng of norfloxacin accumulated per mg dry cell weight.

### EtBr accumulation assay

EtBr accumulation and efflux were performed using a semi-automated fluorometric method.^14, 56, 57^ Briefly, for accumulation assays, mid-log phase. WT *smegmatis* and Nor^r^ mutant cells were centrifuged at 13000 rpm for 3 min, and the pellet was washed and resuspended in PBS (pH 7.4). EtBr was added to the cellular suspension at a concentration of 3 µg/ml (less than MIC_1/2_). 200 µl of this was added to 96 well plates (black polysorp, Nunc), and the EtBr accumulation was measured by measuring the fluorescence in a microplate reader (spectramax multimode M 5, Molecular devices) at 37°C using 530 nm excitation wavelength and 585 nm as the emission wavelength. For studying the effect of efflux inhibitor carbonyl cyanide m-chlorophenylhydrazone (CCCP) on EtBr accumulation, mid-log phase cells were washed and suspended in PBS containing the desired concentration of CCCP. EtBr was added to the cellular suspension at a final concentration of 3 µg/ml, and 200 µl of this was added to 96 well plates for measuring the EtBr accumulation. Fluorescence data were acquired every 60 seconds for 60 minutes. Accumulation data were plotted as relative RFU where the fluorescence value at each time was subtracted with respect to 0 min value.

### Cell permeability assay (NPN assay)

Bacterial cell permeability was measured by using a fluorescent probe, 1-N-phenylnaphthylamine (NPN). NPN has a weak fluorescence in aqueous solution but turns strongly fluorescent in the phospholipid environment.^55^ Briefly mid-log phase *M. smegmatis* WT and Nor^r^ mutant cells were centrifuged to remove the media and dispersed in 0.5 mM HEPES buffer (pH 7.2). Aliquots of 100 μl of the cell suspension and 100 μ μM) were added to 96 well plates (black polysorp, Nunc) and the fluorescence of NPN was monitored in a microplate reader (spectramax multimode M5, Molecular Devices) at 37°C, using 402 nm and 355 nm as excitation and emission wavelengths, respectively.

### Construction of *M. smegmatis Msmeg*_5659 over-expressing strain

Optimization of primers was done at various temperatures using 500 - 600 ng of gDNA from *M. smegmatis* WT, specific forward and reverse primers (0.5 mM final concentration), 1.5 units DreamTaq DNA polymerase (Thermo Scientific), 1x reaction buffer, 0.2 mM dNTP’s and 1.5 mM MgCl_2_ in a thermal cycler (Hi-MEDIA), followed by PCR clean-up was done. Two steps plasmid and gene of interest digestion was done, followed by gel purification. Restriction digestion was done using specific restriction enzymes BamHI and Hind III (0.5 U final concentration) and 1x cut smart reaction buffer (NEB) incubated in the water bath at 37 °C for 2 hours and 15 minutes respectively followed by heat inactivation for 15-20 minutes. The double-digested plasmid and PCR products were run on 1.2% low melt agarose gel, followed by clean-up using the gel purification kit (Qiagen). Ligation of double digested plasmid and insert (PCR purified) was carried out using 1 unit of T4 DNA ligase (Ferments) and 1x ligase buffer in a ratio of 1:7 at 37 °C for 30 minutes and incubated at -4 °C overnight, afterward stored at -20 °C. For competent cell preparation, *E*. *coli* DH5α cells were grown in LB media to an OD_600_ of 0.5, incubated with a 50mM CaCl_2_ solution, followed by re-suspension into 50% ice-cold glycerol. For electro-competent cell preparation, *M. smegmatis* WT cells were grown in M7H9 media to an OD_600_ of 0.5, followed by washing with 10 % ice-cold glycerol. Next, the cells were resuspended into 10 % glycerol (ice cold), i.e., one-fifth of the original volume and stored at -80°C in 60µL aliquots. The transformation was done through electroporation (0.1 cm cuvette) with a final concentration of 1µg of ligated product in 60µL competent cells using the capacitance of 2.5 kV. For confirmation of clone, plasmid (i.e., construct) was isolated from the clone, followed by PCR amplification with specific primers (Table S2).

### DNA isolation, PCR and DNA sequencing

Mid-log phase cells of *M. smegmatis* and *M*. *smegmatis* (pSTKiT) were used for the isolation of genomic DNA. Genomic DNA isolation protocol was optimized in our laboratory based on a previously modified method for mycobacteria.^58^ Briefly, the cells were suspended in 400 µl of nuclease-free water, and the suspension was heat-inactivated at 65 °C for 30 min. Cell wall disruption was performed by mechanical bead-beating using 0.1 mm zirconium beads (Unigenetics). Further, lysis and digestion were achieved by adding lysozyme (30 µl from100 mg/ml stock) followed by incubation at 37 °C for 1 h. To this 70 µl of 10 % SDS and 10 µl of proteinase K (20 mg/ml stock) was added and the resulting suspension was gently vortexed and incubated for 15 min at 65 °C. 100 µl NaCl (5 M) and 100 µl CTAB/NaCl solution (which was pre-warmed at 65 °C) were then added to the suspension. Finally, the DNA was extracted using phenol-chloroform extraction, precipitated with isopropanol, washed with 75 % ethanol and dissolved in autoclaved water. DNA concentration and purity were measured by nanophotometer (Implen). Further, the quality of DNA was also monitored by 1.2% agarose gel electrophoresis using ethidium bromide staining.

Specific genes from and Nor^r^ mutant were sequenced for identifying mutations in drug target and transport-related genes. Primers and their corresponding gene sequences are listed in Table S1. PCR amplification was done by using a standardized protocol. In a typical PCR reaction setup, 100-200 ng of DNA, gene-specific forward and reverse primers (1 mM final concentration), and 5 % DMSO was added to the optimized PCR master mix containing DreamTaq DNA polymerase (Thermo scientific), reaction buffer, dNTPs and MgCl_2_. The PCR reaction was performed in a thermal cycler (BIO-RAD) using optimized thermal cycling conditions. PCR products were sequenced with an automatic DNA sequencer (Applied Biosystems 3730 DNA analyzer) by Eurofins Genomics India Pvt Ltd. DNA sequencing was performed for 2-4 biological replicates.

### Whole Genome Sequencing

Genomic DNA (gDNA) extraction was done using NaCl method to generate fragments of approximately 200-500bp, the size distribution of each fragment was checked using high sensitive DNA kit. Further, the library was prepared using the Truseq Nano DNA library kit for WT gDNA and the NEXTFlex DNA sequencing kit for the Norr gDNA. Sequencing data had been generated on Illumina platform, using paired-end sequencing with a read length of 150 bp and average covergae of 100X. Analysis of the raw sequencing data was performed using the Galaxy web platform at usegalaxy.org.^59^ First, quality check was performed using FastQC. Further, Bowtie 2 aligner was used for aligning the raw reads of both the strains of *M.smegmatis* with the reference genome (Accession number NC_008596), respectively.^60^ Post alignments, Samtools/mpileup was used to sort the bam files.^61^ After the reads were aligned and sorted, the sam files were scanned using Varscan to filter out the variants.^62^ The variants with minimum depth of 50 and frequency greater than 80% were taken into consideration. Further, the variants from both the sam files were mapped using R programming^63^, to bring out the mutations, which are present in the mutant and not in wild type.

### RNA isolation

*M. smegmatis* WT and Nor^r^ mutant cells were grown till they reached OD_600_ of 0.5. A 12% phenol-ethanol mixture was added to the cells to avoid RNA degradation. Cells were centrifuged at 10000 rpm for 10 min. The supernatant was discarded, and the pellet was stored at -80°C until extraction. RNA was extracted using TRI reagent (Sigma). The cell pellet was resuspended in 1 ml of TRI reagent. The cell lysis was performed by mechanical bead-beating using 0.1 mm zirconium beads (Unigenetics) in a mini bead-beater (Unigenetics) for 1.5 min (30-sec pulse). The cell lysate was centrifuged at 12000 rpm for 10 min at 4°C, and the supernatant was transferred to a fresh tube. Phase separation was performed by adding 200 µl of chloroform per ml of TRI reagent (samples were vortexed vigorously for 15 sec; allowed to stand for 10 min at room temperature and centrifuged at 12000 rpm for 15 min at 4°C). The mixture separated into three phases. The upper aqueous phase was transferred to a fresh tube, and an equal volume of 2-propanol was added to precipitate the RNA in the form of a pellet. Samples were centrifuged, and the RNA pellet was washed three times, with 75 % ethanol. The pellet was air-dried for 5-10 min and dispersed in nuclease-free water. Isolated RNA was DNase treated with Turbo DNase (Ambion) to get rid of any trace DNA contamination. RNA quantity and quality were checked by measuring absorbance ratio (A260/A280) using a nanophotometer (Implen), and the integrity of RNA (23S and 16S bands) was confirmed on a 1.2% denaturing agarose gel.

### Reverse transcription

First-strand cDNA was synthesized by reverse transcription from RNA using the RevertAid^TM^ H Minus Reverse Transcriptase kit (Fermentas) as per the manufacturer’s instructions. Briefly, four µg of total RNA was added to 100 pmol of random primers and water to make volume up to 13 µl. Mycobacterial RNA being GC-rich, the mixture is heated at 65°C for 5 min followed by immediate chilling on ice. To this mixture, 4 µl of 5X reaction buffer, 2 µl of dNTP MIX (10 mM each) and 1 µl of Moloney Murine Leukemia Virus (M-MulV) reverse transcriptase III was added such that total volume becomes 20 µl. The reaction was performed in a thermal cycler (BIO-RAD) (25°C for 10 min, followed by increasing the temperature to 45°C for 60 min, followed by termination at 70 °C for 10 min). A no reverse transcriptase (NRT) control was also set up where the mixture did not contain the reverse transcriptase enzyme to quantify gDNA contamination.

### Quantitative real-time PCR (qRT PCR)

Quantitative real-time PCR (qRT-PCR) was performed on a iQ5 cycle (BIO-RAD) for gene expression analysis. 100 ng of cDNA was used as a template for qRT-PCR. The reaction was provided with 0.5 µM gene-specific primers and SYBR green (iQ5 SYBR Supermix-BIO-RAD). The lists of primers used in this study are given in Table S2. These primers were designed using Primer 3 (http://bioinfo.ut.ee/primer3-0.4.0/). The PCR cycle consisted of incubation at 95°C for 3 min, followed by 40 cycles of denaturation at 95°C for 1 min, annealing for 1 min (see tables S3 for optimized temperature specific for each primer) and extension at 75°C for 1 min. At the end of each PCR run, a melt curve analysis was performed for confirming the amplification of a single product. *sig*A gene was used as the housekeeping gene. Relative gene expression values in terms of log_2_fold change were calculated using the 2^-ΔΔ^ where all the values are normalized to a housekeeping gene (*sig*A RNA) as shown below:

ΔC_T_^1^ = C_T_ (GOI-treated) –C_T_ (HK-treated)

ΔC_T_^2^ = C_T_ (GOI-control) – C_T_ (HK-control)

ΔΔC_T_ = ΔC_T_^1^ – ΔC_T_^2^

A Fold Change of Expression = 2^-ΔΔCT^_, where_ GOI is a gene of interest, HK is a housekeeping gene, and C_T_ is the threshold cycle.

## ACKNOWLEDGMENTS

We acknowledge Science and Engineering Research Board (SERB), Department of Science & Technology (DST), Government of India for funding through a project (sanction letter No.: EMR/2016/007667). We would like to acknowledge Sophisticated Analytical Instrumental Facility (SAIF), IIT Bombay and Industrial Research and Consultancy Centre (IRCC), IIT Bombay for providing SEM, Cryo-FEG-SEM, and HR-LCMS facilities. DR acknowledges SERB, DST, Govt. of India for her fellowship through a project (sanction letter No. SB/S3/CE/059/2014). We also acknowledge the help of Monali Praharaj for initial experiments on the Nor^r^ mutant and Prof. Markus A. Seeger from University of Zurich, Zurich, Switzerland for generously providing us the deletion mutant of *Msmeg*_5659 used in this study.

## Supporting information

- Table S1: List of Primers and optimized temperatures for sequencing and transcript analysis.
- Table S2: List of Primers and restriction sites used for cloning.
- Table S3: MIC profile against the various drug of Msmeg_5659 with inducer concentration 40ng/ml overtime.
- Table S4: Effect of knockout of efflux pump gene *Msmeg*_5659 on drug susceptibility.

