## Supplemental file for "Laboratory evolution of *Mycobacterium smegmatis* in the presence of a fluoroquinolone leads to extreme drug resistance phenotype through the over-expression of *Msmeg_5659-61 efflux pump.*"

**Supplementary information**

### Table S1: List of Primers and optimized temperatures for sequencing and transcript analysis

| **Seq. Name** | **Tm Opt. (˚C)** | **Sense Primer** | **Anti-sense Primer** |
| --- | --- | --- | --- |
| **Sequencing Primers** | | | |
| ***gyrA*** | 54 | TTTAACCCGCGATCCGACAA | CGGCTGTCCTCCTCGATCTC |
| ***gyrB*** | 54 | CAAGGATCCCAACCTCACCG | CGTTGCGCGTGATGAAGCTA |
| ***rpoB*** | 54 | GCATGACCACCCAGGACG | GCAGGCGGTAGGACTGAC |
| ***katG1*** | 54 | GCGAAAGCACGAACTTCCC | AAGCGGGTTCTCCAGGTCA |
| ***katG2*** | 54 | GGCCGACCTGATCGTGTATG | CCTTCCACGCGGTCGAAAC |
| ***katG3*** | 54 | GGCTGTGGCAGGACAACAT | GGTGGTTGTGTGTCAGGCG |
| ***lfrA1*** | 64 | TTATGAGGCAGCCAAGGACG | GCCGATGATCGACAGGAAGT |
| ***lfrA2*** | 64 | AGCATCGTGCTGTCGTTCA | CTACGTCTACGACCCCGAGC |
| ***lfrR*** | 64 | CCTCCTCCGCCATCTCCG | CGATGCAGGTGGACATGAGA |
| **qRT Primers** | | | |
| ***SigA*** | 52 | GTGCATGTCAAACCCAGGTAAGG | GGGATCCGTGCCGTAGCTAAC |
| ***MSMEG__6225 (lfrA)*** | 60 | ATCTCGCAGCACCTTCAGTT | CGGAACAACAGGATCATCAG |
| ***MSMEG_2619 (efpA)*** | 62 | TCGGATTCATCCCGTTCGTG | GTGCAGTGTCGAACCGTAGA |
| ***MSMEG_3069 (p55)*** | 55 | ATCTGCTCGGCTACATCGC | GATCACCAGCATCGTCAGGT |
| ***MSMEG_5659*** | 53 | AGAAGGTGCTGCTGGAGTTG | CTCCAGCATGTCCTGGATGG |
| ***MSMEG_4700*** | 58 | ACCGCTACGAGGAGATGGT | GTTGAAGCCCTTGTCGAGGT |
| ***MSMEG_3763*** | 57 | TGGTTGCTGGGGTTCCATAC | GATGACCAGCGGCATGAACT |
| ***MSMEG_0250*** | 58 | TCTACGACGTGGATTCCGGC | TTGTCGTAGATCGTGCCCGC |
| ***MSMEG_5009*** | 66 | ACGCTCAAGCTCGACTACAC | GAAGCGGCCCGAGATGTAAT |
| ***MSMEG_0241*** | 58 | AGAAGGTGCTGCTGGAGTTG | CTCCAGCATGTCCTGGATGG |
| ***pstB*** | 55 | ACGCTCAAGCTCGACTACAC | GAAGCGGCCCGAGATGTAAT |
| ***MSMEG_5003*** | 58 | TTCGGTGCGTTCTTCGATCC | TGTAGGCGAGGTTGAACACC |

### Table S2: List of Primers and restriction sites used for cloning

| **Constructs** | | **Primers** | **Enzymes** |
| --- | --- | --- | --- |
| *Msmeg*_5659_OE/i_ | *Msmeg*_5659 | 5’ GAAGAAAATCAGGCCAGCGAG 3’ | BamHI |
|  |  | 5’ AGTGGGAGGACGACGAGG 3’ | XbaI |
|  | *Msmeg*_5660 | 5’GAATATTGAGTGGTTGTGACTGAACC 3’ | XbaI |
|  |  | 5’ GGGTCCTGCTCCTCCTGC 3’ | EcoRI |
|  | *Msmeg*_5661 | 5’ GTGAGCAAATGAGTATCGAGACC 3’ | EcoRI |
|  |  | 5’ GGCAACTGAATAACCTCACGC 3’ | HindIII |

### Table S3: MIC of various drugs against Msmeg_5659_i_ strain. The inducer, tetracycline was used at a concentration of 40ng/ml for varying induction times as shown below.

| Drug class | Drug | *Msmeg*_5659_i_ (Inducer concentration 40ng/ml) | | | | |
| --- | --- | --- | --- | --- | --- | --- |
|  |  | 0H | 1H | 2H | 5H | 14H |
| FQs | Norfloxacin | 2 | 8 | 8 | 8 | 16 |
|  | Ofloxacin | 0.6 | 2.4 | 2.4 | >2.4 | 4.8 |
|  | Ciprofloxacin | 0.4 | >0.8 | >0.8 | >0.8 | 1.6 |
|  | Moxifloxacin | 0.08 | 0.16 | 0.16 | 0.16 | 0.16 |
| 1^st^ line | Rifampicin | 32 | 64 | 64 | 512 | 512 |
|  | Isoniazid* | 32 | 32 | 64 | 128 | 256 |
|  | Streptomycin | 1 | >4 | 4 | >4 | 16 |
| 2^nd^ line | Amikacin | 0.5 | >2 | 1 | >2 | 4 |
| Aromatic | EtBr | 7 | 14 | 14 | 14 | 14 |

**Table S4: Effect of knockout of efflux pump gene *Msmeg*_5659 on drug susceptibility**

| Drug class | Drug | WT | Δ Msmeg_5659 | Msmeg_5659_OE_ |
| --- | --- | --- | --- | --- |
| FQs | Norfloxacin | 2 | 2 | 16 |
|  | Ofloxacin | 0.6 | 0.6 | 2.4 |
|  | Ciprofloxacin | 0.4 | 0.4 | 3.2 |
|  | Moxifloxacin | 0.08 | 0.08 | 0.16 |
| 1^st^ line | Rifampicin | 32 | 32 | 128 |
|  | Isoniazid* | 32 | 32 | 256 |
|  | Streptomycin | 1 | 1 | 2 |
| 2^nd^ line | Amikacin | 0.5 | 0.5 | 4 |
| Aromatic | EtBr | 7 | 7 | 14 |
